# Immune signatures correlate with L1 retrotransposition in gastrointestinal cancers

**DOI:** 10.1101/216051

**Authors:** Hyunchul Jung, Jung Kyoon Choi, Eunjung Alice Lee

## Abstract

Long interspersed nuclear element-1 (L1) retrotransposons are normally suppressed in somatic tissues mainly by DNA methylation and antiviral defense. However, L1s can be desuppressed in cancers to act as insertional mutagens and cause genomic instability by creating DNA double strand breaks and chromosomal rearrangements. Whereas the frequency of somatic L1 insertions varies greatly among individual tumors, much remains to be learned about underlying genetic, cellular, or environmental factors. Here, our pan-gastrointestinal cancer genome analyses for stomach, colorectal, and esophageal tumors identified multiple correlates of L1 activity. Clinical indicators of tumor progression, such as tumor grade and patient age, showed positive association. Potential L1 expression suppressors such as *TP53* and *DNMT1*, a DNA methyltransferase, were inactivated in tumors with frequent L1 insertions. Importantly, tumors with high immune activity, for example, due to viral infection or high tumor-antigen load, tended to carry a low number of L1 insertions in their genomes with high expression levels of L1 suppressors such as *APOBEC3s* and *SAMHD1*. Our analysis of the transcriptional effects of intragenic retrotransposon insertions demonstrated an increased risk of gene disruption in retrotransposition-prone cancers. In particular, we found a splicing-disrupting L1 insertion in an exon of *MOV10*, a key L1 suppressor, which caused exon skipping with evidence of nonsense-mediated decay in a tumor with a high L1 insertion load. Our results indicate that cancer immunity may contribute to genome stability by suppressing L1 retrotransposition particularly in gastrointestinal cancers.

## INTRODUCTION

Frequent desuppression and retrotransposition of the long interspersed nuclear element-1 (LINE-1 or L1) have been reported in multiple cancer types (Lee et al. 2012; Solyom et al. 2012; Helman et al. 2014; Tubio et al. 2014). Notably, gastrointestinal cancers, including esophageal (Doucet-O’Hare et al. 2015; Secrier et al. 2016), gastric (Ewing et al. 2015), and colorectal cancers (Lee et al. 2012; Solyom et al. 2012), reportedly carry extensive somatic L1 insertions. The rate of L1 insertions varies substantially among individual tumors, ranging from a few to hundreds. Clinical and molecular factors identified in association with L1 insertions include patient age in colorectal cancer (Solyom et al. 2012), patient survival in pancreatic cancer (Rodic et al. 2015), and *TP53* mutations in head and neck cancer (Helman et al. 2014). However, further investigation is needed on the mechanisms underlying these associations. Furthermore, previous studies may have been limited in their ability to detect other factors, especially those related to major L1 suppression mechanisms, namely DNA methylation and antiviral defense, due to small sample sizes and/or lack of matched expression profiles.

L1 insertions are able to disrupt target gene function by interrupting protein-coding sequences or altering mRNA splicing and expression. Intragenic somatic L1 insertions previously identified in cancer genomes were depleted in exons and mostly located in introns, generally decreasing target gene expression (Lee et al. 2012; Helman et al. 2014) with some exceptions (Helman et al. 2014). On the other hand, Tubio et al. analyzed 24 expression profiles from TCGA lung and colon cancer samples and reported no evidence of altered gene expression and aberrant transcripts caused by somatic L1 insertions (Tubio et al. 2014). Although aberrant splicing is a major pathogenic mechanism of retrotransposon insertions causing Mendelian disorders and hereditary cancers (Hancks and Kazazian 2016), to our knowledge, no somatic L1 insertions have been reported in association with splicing alterations in sporadic human cancers.

Here, we analyzed whole genome sequencing data for which somatic retrotransposition had not previously been investigated, and which were obtained from cancer patients of three gastrointestinal cancer types using an improved version of *Tea* (Transposable Element Analyzer) (Lee et al. 2012). Among other findings, our analysis identified cancer immunity as the most prominent variable that explained variation in retrotransposition rates among individual tumors. We identified exon skipping caused by somatic retrotransposon insertions and found a significant downregulation of genes with somatic L1 insertions, corroborating the high gene disrupting potential of somatic retrotransposition in cancer.

## RESULTS

### Highly variable frequency of somatic L1 insertions in gastrointestinal cancers

We applied *Tea* (Transposable Element Analyzer) (Lee et al. 2012) with improved 3’ transduction (*i.e.*, mobilization of unique non-L1 DNA downstream of the L1) detection to the whole-genome sequencing data of tumor and blood samples from a total of 189 gastrointestinal cancer patients across three cancer types: 95 stomach (40 TCGA and 55 non-TCGA (Wang et al. 2014)), 62 TCGA colorectal, and 32 esophageal (19 TCGA and 13 non-TCGA (Dulak et al. 2013)) cancer patients. We detected 3,885 somatic L1 insertions with target site duplication (TSD) and polyA tails (**Supplemental Table S1** and **S2**), the two signatures for target-primed reversed transcription (TPRT)-mediated retrotransposition. While the insertion frequency varied greatly, most (89%) samples carried at least one insertion with the average number of insertions at 21 (Fig. 1a and **Supplemental Table S3**), thereby confirming previous findings that gastrointestinal cancers are highly susceptible to somatic L1 retrotransposition (Burns 2017). Out of 137 insertions with 3’ transduction, more than a half (56%) were derived from two germline L1s on chromosome X and 22 (Xp22.2 and 22q12.1) (Fig. 1b and **Supplemental Table S2),** consistent with a previous finding that a handful of source L1s generated most 3’ transductions in cancers (Tubio et al. 2014).

1,192 (31%) of the 3,885 L1 insertions were found in gene bodies—mostly in introns (29%, Fig. 1c). A total of 210 genes, including known (*LRP1B* and *PTPRT*) and putative cancer driver genes were affected by somatic L1 insertions in multiple cancer samples (Fig. 1d). For example, *ROBO1*, an emerging tumor suppressor (Gara et al. 2015; Huang et al. 2015), had at least one somatic L1 insertion in each of seven cancer samples (two stomach and five colorectal samples). *PARK2*, a master regulator of G1/S cyclins that is frequently deleted in cancers (Gong et al. 2014), was marked by one insertion in each of six samples (one stomach, three colorectal, and two esophageal samples). Genes with recurrent somatic L1 insertions (Fig. 1d), including *ROBO1, PTPRT, GRID2, CDH8, CDH12, CDH13, PTPRM, ROBO2,* were enriched for brain development and function including axon guidance, neuron differentiation, and synaptic function (**Supplemental Table S4**). However, this could be attributed to the fact that neuronal genes tend to be long (Zylka et al. 2015). Indeed, no significant enrichment was found after adjusting for gene size, suggesting the overall absence of positive selection of cancer cells with somatic L1 insertions.

**Figure 1.**
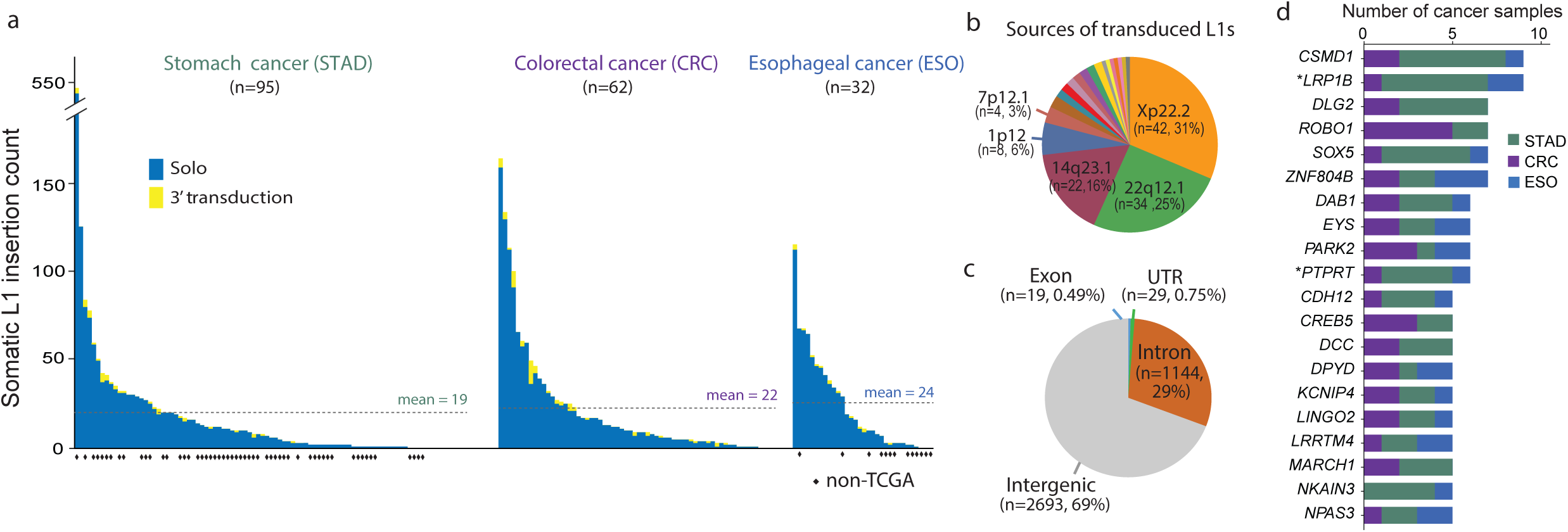
Landscape of somatic L1 insertions in gastrointestinal cancers. (a) Frequency of somatic L1 insertions across three cancer types. The dotted line denotes the average insertion count for each cancer type. (b) Source L1 elements of somatic L1 insertions with 3’ transduction. (c) Genomic distribution of somatic L1 insertions. (d) Genes with recurrent somatic L1 insertions. Genes with somatic L1 insertions in more than four cancer patients are shown. Known cancer genes reported in the COSMIC Cancer Gene Census database v82 are marked with an asterisk.

### Cancer immune activity negatively correlated with somatic L1 retrotransposition

We then wanted to understand mechanisms underlying the variable frequency of L1 insertions in cancers. First, we examined the association of L1 insertions with molecular markers or clinical traits. We observed a significant association of *TP53* mutation status with the L1 insertion rate. Somatic L1 insertions were more frequent in tumors with *TP53* mutations than those with wild-type *TP53* (*P* = 0.004, Fig. 2a). This corroborates the recent implication of *TP*53 in restraining L1 transcription (Wylie et al. 2016). When we examined whether any aberration in DNA repair pathways could be associated with L1 retrotransposition, only the p53 repair pathway showed a significant association (*P* = 6.3 × 10^−3^, see Methods for details). Regarding associations with tumor grade and patient age, more frequent somatic L1 insertions were observed in cancers at an advanced stage (*P* = 0.043, Fig. 2b) and in older stomach cancer patient samples (*P* = 0.054, Fig. 2c). We found a positive correlation between L1 insertion frequency and the expression level of L1 itself (*P* = 0.0052, Fig. 2d) and a negative correlation with the expression level of *DNMT1,* a DNA methyltransferase (r = −0.31, *P* = *7.2 × 10^−4^*). These associations suggest that aberrant L1 transcription potentially induced by DNA methylation loss and mutations in L1 transcription suppressors is a prerequisite to frequent retrotransposition in cancer.

**Figure 2.**
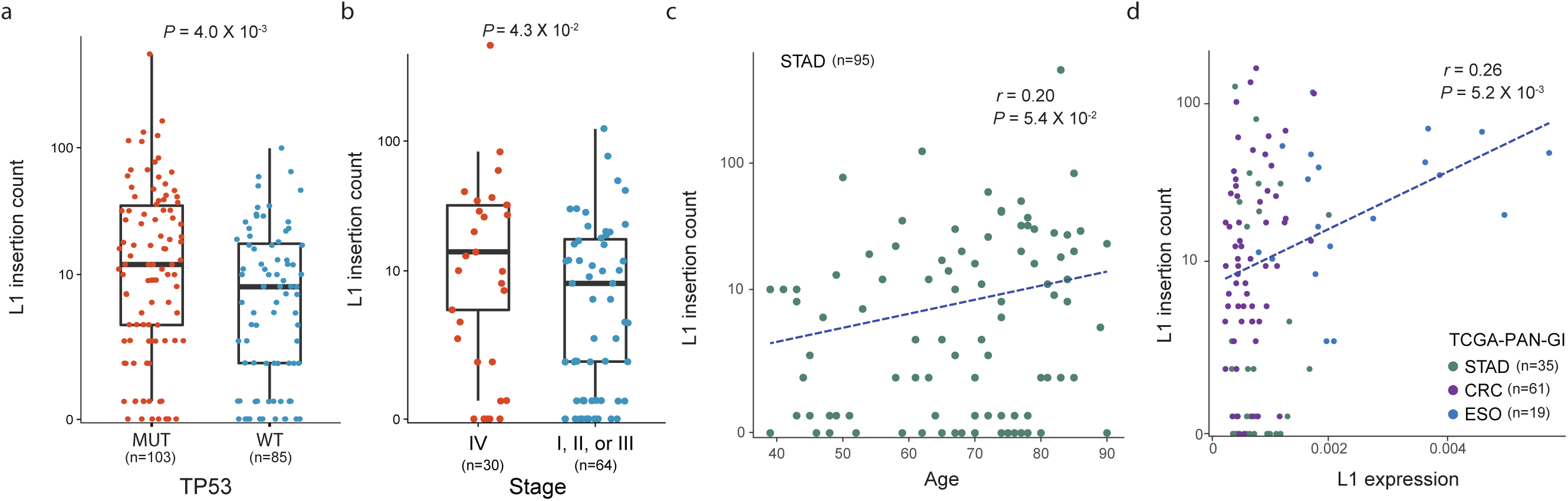
Factors correlated with the frequency of somatic L1 insertions. (**a**) Somatic L1 insertion counts in cancer samples with mutations (MUT, red dots) and without mutations (WT, blue dots) in *TP53* are shown in boxplots. (**b**) Somatic L1 insertion counts in stomach cancer samples at stage 4 (red dots) and at earlier stages (stage 1-3, blue dots) are shown in boxplots. (**c**) Correlation between the age of cancer patients at diagnosis (x-axis) and somatic L1 insertion counts (y-axis) in stomach cancer. (**d**) Correlation between L1 expression (x-axis) and somatic L1 insertion counts (y-axis).

We then performed a more systematic transcriptome analysis by measuring the transcriptional activity of 1,789 pathways from the Reactome database (Milacic et al. 2012; Fabregat et al. 2016) in 112 TCGA cancer samples with RNA-seq profiles, using the single-sample GSEA (ssGSEA) method (Subramanian et al. 2005). A total of 50 and 97 pathways showed positive and negative correlations, respectively, with L1 insertion counts (Fig. 3a **and Supplemental Table S5**, FDR < 0.05). Notably, 49 out of 176 (28%) immune pathways showed significant negative correlations. For example, cancers with active interferon alpha/beta signaling carried fewer somatic L1 insertions (Fig. 3b, r = −0.42, *P* = 4.6 × 10^−6^). To further test the relationship between L1 insertions and immune activity, we used fifteen annotated sets of immune genes (Breuer et al. 2013; Rooney et al. 2015). Our analysis showed consistent negative correlations, especially in stomach cancer (Fig. 3c), thereby supporting a robust immunological association of L1 retrotransposition. Finally, we tested 13 genes that are known to act as L1 inhibitors (Goodier 2016). We found significant negative correlations for *AICIDA*, *SAMHD1*, and *APOBEC3C/D/F/H* (Fig 3d). Since several L1 inhibitors, such as *MOV10* and *APOBEC3* family proteins, are known to be activated by type I interferons (IFNs) (Yu et al. 2015), it is likely that active interferon alpha/beta signaling may suppress L1 retrotransposition by activating L1 suppressors.

**Figure 3.**
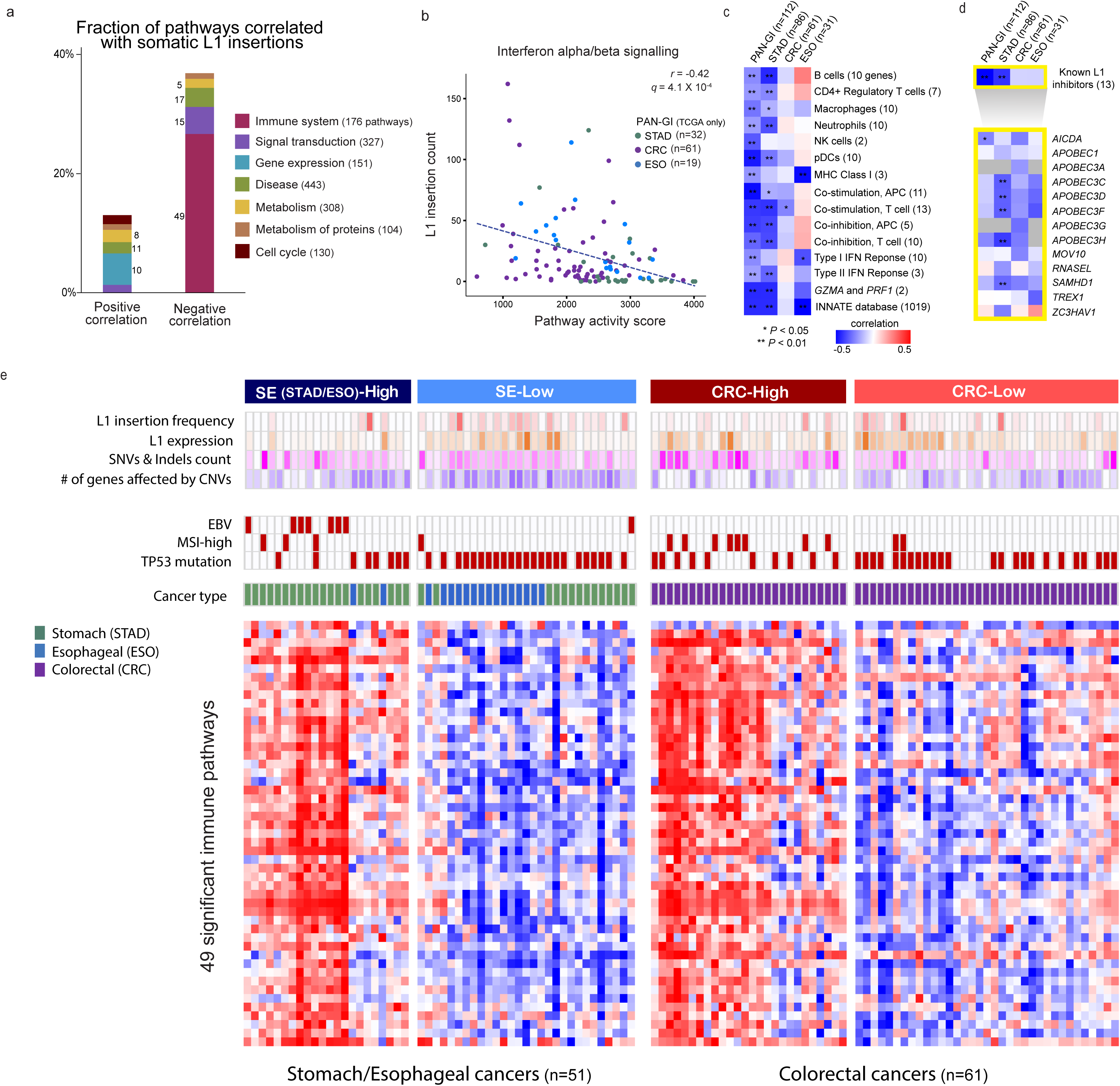
Immune activity associated with somatic L1 retrotransposition. (**a**) Reactome pathways for which activity correlates with somatic L1 insertion frequency. For each category of pathway, the percentage of pathways showing a significant positive or negative correlation between pathway activity and somatic L1 insertion frequency (FDR < 5%) is shown in a stacked bar. The number of significantly correlated pathways for each category is shown next to the stacked bar. The number of member pathways for each category is shown in parentheses, and only the categories with more than 100 member pathways are shown. (**b**) Negative correlation between the activity of Interferon alpha/beta signaling pathway and the frequency of somatic L1 insertions. (**c**) Negative correlation between the activity of various immune gene sets and the frequency of somatic L1 insertions. Each cell in the heatmap shows a color-scaled Spearman correlation coefficient between the activity of each gene set (row) and the frequency of somatic L1 insertions in cancer samples from each cancer type (column). The color scale indicates the degree of correlation. Correlations with *P* < 0.05 and *P* < 0.01 are marked by single and double asterisks, respectively. (d) Correlation between the expression level of known L1 inhibitors and the frequency of somatic L1 insertions. Each cell in the heatmap shows the correlation coefficient between the expression level of an individual L1 inhibitor gene (row) and the frequency of somatic L1 insertions in each cancer type (column). (**e**) Cancer subgroups with distinctive immune activity signatures. The heatmap represents the activity of 49 significant Reactome immune-related pathways (row) in 112 cancer samples (columns). Unsupervised clustering identifies cancer subgroups according to immune activity: stomach/esophageal (SE)-High, SE-Low, colorectal (CRC)-High, and CRC-Low. Clinical and molecular features of each cancer sample are marked on the top of the heatmap. Higher L1 insertion frequency, L1 expression, somatic SNV/indel counts, and counts of genes with somatic copy number aberrations are marked in darker red, orange, pink, and purple, respectively. The color scale was normalized separately for stomach-esophageal cancer and colorectal cancer samples. Cancer samples with EBV infection, MSI-high phenotype, and non-synonymous mutations in *TP53* are marked with filled boxes.

### Characterization of cancer subgroups with differential immune activity

Based on the 49 L1-associated immune pathways in 112 cancer samples, we identified two distinct cancer subgroups for stomach/esophageal cancer (SE; SE-High and SE-Low) and colorectal cancer (CRC; CRC-High and CRC-Low) that differed in their immune signatures (Fig. 3e). Somatic L1 insertions were significantly less frequent in the high immune activity subgroups (Fig. 3e and Fig 4a, *P* = 7.3 × 10^−5^ and *P* = 2.1 × 10^−2^ for SE and CRC, respectively). The expression levels of two cytotoxic T cell effector genes (*GZMA* and *PRF1*) (Rooney et al. 2015) indicated consistent and significant differential cytolytic activity between the subgroups (**Supplemental Fig. S1a**, *P* = 6.4 × 10^−7^ and *P* = 2.9 × 10^−4^ for SE and CRC, respectively). Interestingly, adaptive immune pathways showed more differential activities between the subgroups than innate immune and cytokine signaling pathways (**Supplemental Fig. S2** and **Supplemental Table S6**). For example, ‘*immunoregulatory interactions between a lymphoid and a non-lymphoid cell’* was one of the most differential pathways (*FDR* = 6.9 × 10^−7^ and 8.5 × 10^−7^ for SE-High *vs* SE-Low and CRC-High *vs* CRC-Low, respectively). In fact, this pathway includes multiple receptors and cell adhesion molecules, such as *KIR*s and *LLIR*s, that play a key role in regulating immune cell responses to tumor-antigens. *PD-1 signaling* is another adaptive immune pathway with significant differential activity between the subgroups (*FDR* = 1.5 × 10^−6^ and 6.6 × 10^−7^ for SE-High *vs* SE-Low and CRC-High *vs* CRC-Low, respectively).

**Figure 4.**
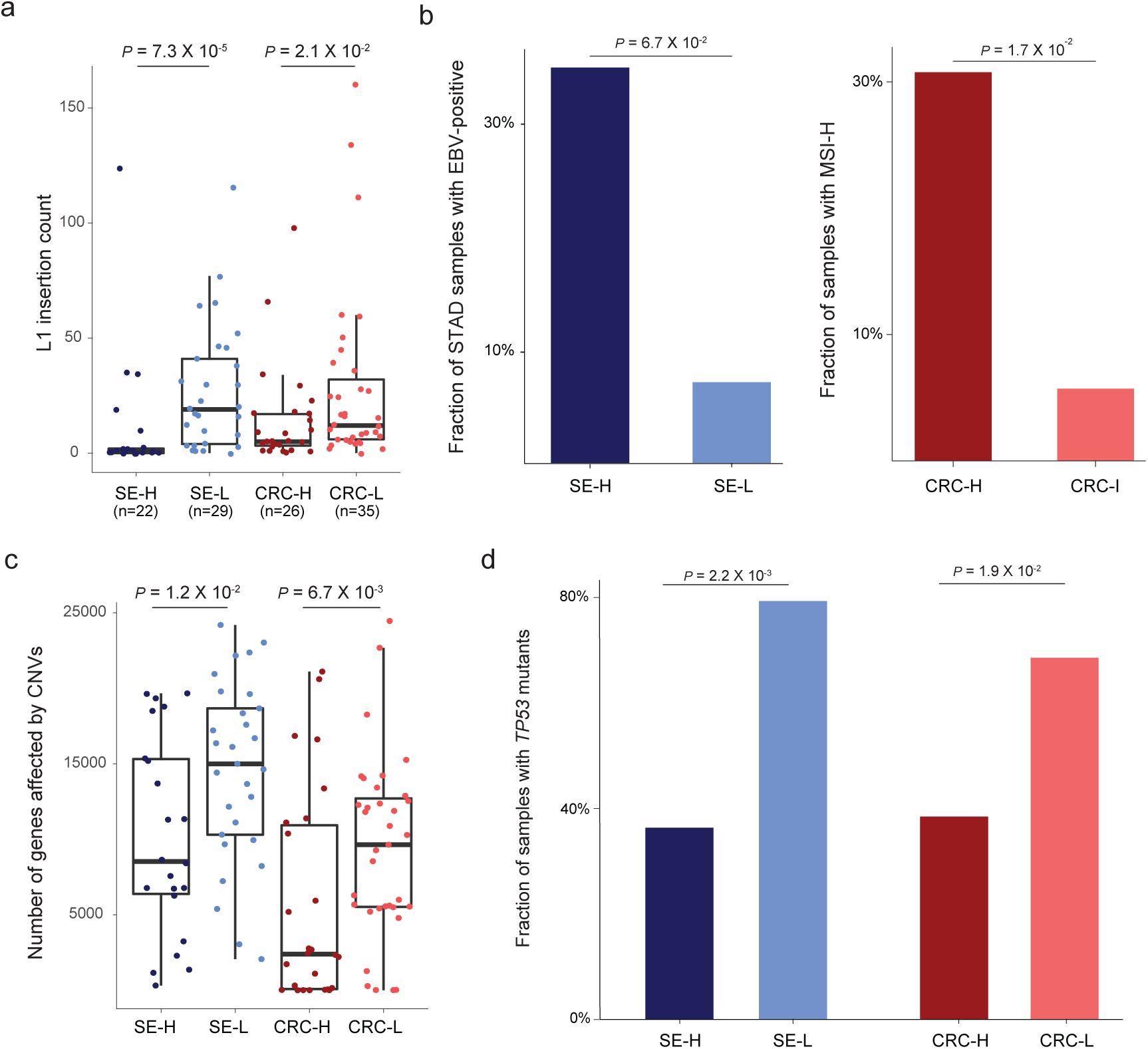
Cancer immunity and genomic instability correlated with somatic L1 retrotransposition. (a) Frequent somatic L1 insertions in low immune activity subgroups. (b) EBV infection and tumor antigens driving high immune response in stomach and colorectal cancer, respectively. The percentage of stomach cancer samples infected by EBV and the percentage of colorectal cancer samples with high microsatellite instability are shown for each immune activity subgroup. (c) Frequent copy number alteration in low immune activity subgroups. The number of genes with somatic copy number aberration in each cancer sample is shown with a colored dot. (d) Frequent *TP53* mutations in low immune activity subgroups. The percentage of caner samples with non-synonymous mutations in *TP53* is shown for each immune activity subgroup.

We next investigated features that could be a potential determinant of immune activity. First, Epstein–Barr virus (EBV) infection was associated with immune activity in stomach and esophageal cancer (left in Fig 4b, *P = 0.067*) whereas microsatellite instability (MSI) and SNV/Indel counts were related to immune activity in colorectal cancer (right in Fig. 4b, *P* = 0.017 for MSI; **Supplemental Fig. S1b**, *P* = 0.0017 for SNV/Indel counts). These suggest that exogenous pathogens such as EBV, and endogenous tumor neoantigens, for example, induced by MSI, may drive high immune activity in gastrointestinal cancer. A recent study reported reduced cytotoxic leukocyte infiltration in highly aneuploid tumors (Davoli et al. 2017). Indeed, the degree of copy number changes significantly differed according to immune activity (Fig. 4c), suggesting the copy number alteration of immune-related genes as another determinant of immune activity (Gao et al. 2016). We also note that mutant *TP53* was associated with low immune activity (Fig. 4d), consistent with previous findings that *TP53* dysfunction leads to immunosuppression (Rooney et al. 2015; Cui and Guo 2016). This result highlights the crucial role of *TP53* in restricting retrotransposons as a guardian of L1 expression (Fig. 2a and **Supplemental Fig. S1c**) and immune integrity (Fig. 4d).

Some cancers may exhibit weak immune activity due to the immune suppressive effect of IFN-signaling, possibly triggered by persistent L1 expression. INF signaling is known to be immune stimulatory, but its persistent activation can trigger immune suppression, especially in the presence of chronic viral infection (Minn and Wherry 2016). Thus, elevated L1 expression mainly caused by DNA hypomethylation and/or *TP53* mutations might activate IFN-signaling as an L1 suppression mechanism (Yu et al. 2015); however, its persistent activation might shift its role to immune suppressor, leading to weakened L1 suppression and thus increased L1 retrotransposition.

### L1 insertions disrupting mRNA splicing and expression

Finally, we wanted to examine the effect of intragenic L1 insertions on transcriptional regulation. To this end, we analyzed matched RNA-sequencing data from 112 TCGA cancer samples for which genomes were analyzed for L1 insertions. Briefly, we calculated the ratio of abnormally spliced RNA-seq reads to normally spliced reads near a somatic L1 insertion and evaluated whether the ratio was significantly higher than expected given the ratio distribution estimated from RNA-seq profiles of control samples without the given insertion.

We screened 1192 intragenic L1 insertions with matched RNA-seq profiles and found skipping of exon 20 of *MOV10,* a known L1 suppressor, with a somatic L1 insertion in one esophageal cancer sample (Fig. 5a). The cancer sample carried 65 somatic L1 insertions and belonged to the low immune group. *MOV10* is known to suppress L1 expression and decrease cytoplasmic L1RNPs (Goodier et al. 2012). The exon skipping event caused by the insertion is likely to have disrupted the *MOV10* function through multiple mechanisms, including the creation of premature termination codons (PTC) followed by nonsense-mediated decay (NMD). We hypothesized that if an L1 insertion triggers NMD, we might observe decreased expression of transcripts with the insertion allele. Since it is hard to distinguish RNA-seq reads from an L1 insertion allele, we used heterozygous single-nucleotide variants (SNVs) to assess allele-specific down-regulation of *MOV10*. The analysis revealed a significantly lower expression level of one allele of *MOV10* in the L1 insertion-carrying cancer sample than in cancer samples with no mutations of any type in *MOV10*, including SNVs, CNVs, and DNA methylation (Fig. 5c). Our additional screening of published 282 somatic retrotransposon insertions (Helman et al. 2014) in 69 cancer samples identified another exon skipping event in a lung cancer sample caused by an Alu insertion in exon 4 of *CYR61*, a putative tumor suppressor (Tong et al. 2001) **(Fig. 5b)**.

**Figure 5.**
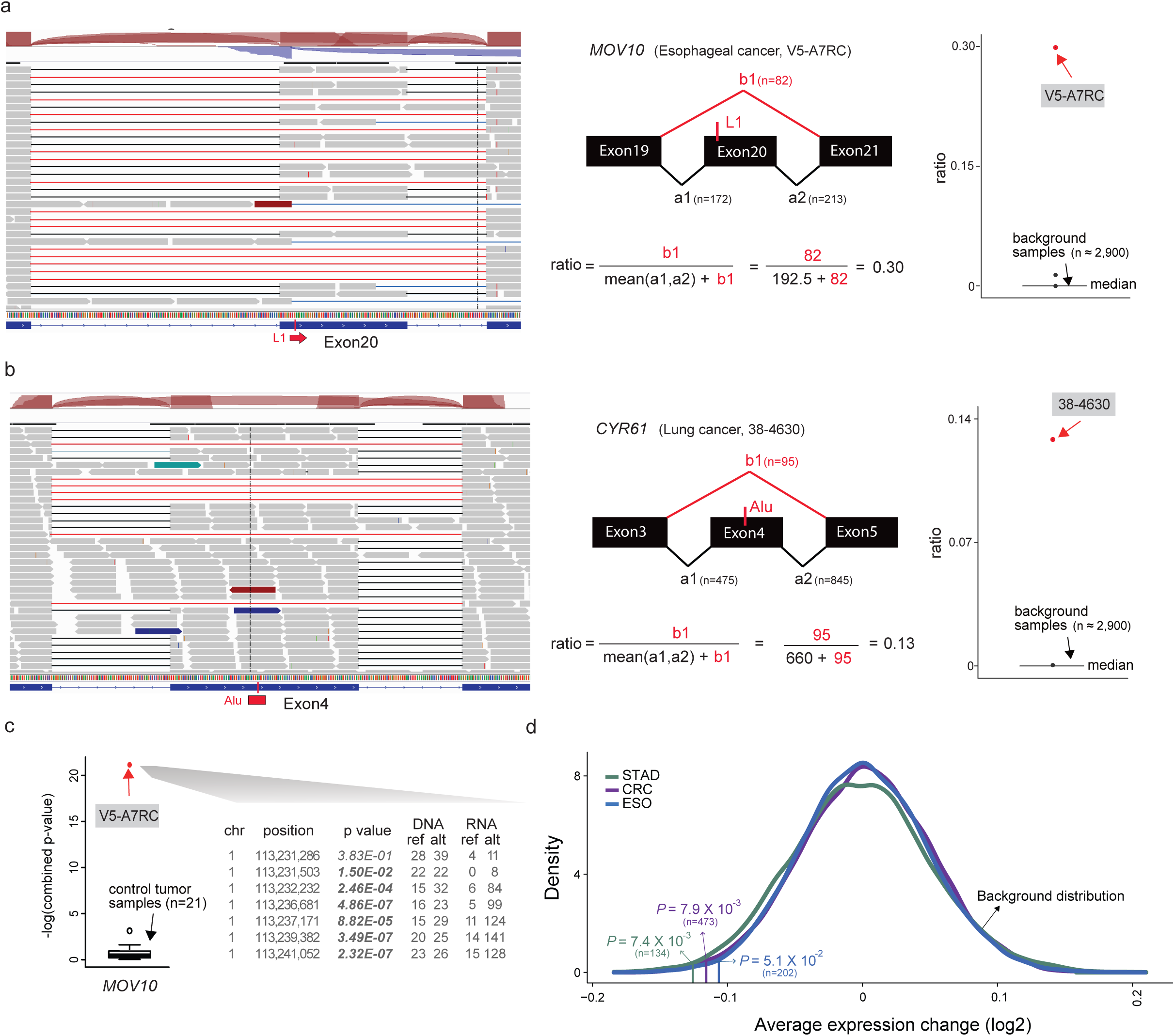
Disruption of mRNA splicing and expression by somatic L1 insertions. (**a**) Skipping of exon 20 in *MOV10* caused by a somatic L1 insertion in TCGA esophageal cancer sample. RNA-seq reads (grey box) from normally spliced transcripts show split mapping between the expected adjacent exons (black lines) whereas reads from transcripts with exon skipping show abnormal split mapping without the exon with the L1 insertion (red lines). Another minor form of abnormal splicing that involves retention of the 19th intron, partial skipping of exon 20, and skipping of exon 21 is also shown with blue lines. The schematic diagram in the middle shows how to calculate the ratio of the abnormally spliced read count (major form) to the total read count around the exon with the L1 insertion. The ratios are also calculated using ~2900 control RNA-seq profiles (cancer samples without any mutation in *MOV10* and normal tissue samples) and serve as a background distribution to assess the significance of the observed ratio from the esophageal cancer sample (red dot). (**b**) Skipping of exon 4 in *CYR61* caused by a somatic Alu insertion. Reads from the 3rd and the 4th introns were often observed in the control RNA-seq profiles, suggesting that they were pre-spliced transcripts not associated with the Alu insertion. (**c**) Increased allelic-bias of *MOV10* in mRNA expression relative to that in DNA sequence in an esophageal cancer with exon skipping caused by a somatic L1 insertion shown in (a). The esophageal cancer sample had seven heterozygous SNVs reported in dbSNP in *MOV10*. For each of the seven SNV loci, the number of reads with a reference allele and an alternate allele was extracted from whole genome sequencing and also RNA sequencing data. Then, the degree of allelic-bias in RNA-sequencing data relative to DNA-sequencing data was tested using Fisher’s test, and loci with significant bias (*P* < 0.05) are marked in bold. The combined P value from seven loci in the esophageal cancer sample (red dot) is shown compared to the distribution of the combined P values from 21 control cancer samples without any mutation and methylation aberration in *MOV10* (black boxplot). (**d**) Decreased expression of genes with somatic L1 insertions. The average difference for each cancer type is marked by a vertical line. The P value of the observed average expression difference was calculated based on a background distribution estimated from random gene sets for each cancer type (colored curved line). The number of genes with somatic L1 insertions for each cancer type is shown in parentheses.

Recent work has reported that L1s tend to insert into lowly expressed genes (Tubio et al. 2014). We thus investigated the expression level of genes with somatic L1 insertions in our data. Out of 809 expressed genes (mean TPM > 0 per cancer type) with one or more somatic L1 insertions, 167 (21%) and 111 (14%) genes expressed at a moderate (TPM 10~100) and high (TPM ≥100) level, respectively (**Supplemental Fig. S3**). This shows that one-third of all intragenic insertions have the potential to alter target gene expression levels. To test this possibility, we next compared the expression level of genes with L1 insertions with the expression level of the same genes in cancer samples without L1 insertions and non-synonymous mutations in the genes. Our analysis revealed that L1 insertions significantly disrupt the expression of target genes in all cancer types, consistent with our previous finding (Lee et al. 2012) (Fig. 5d). The significant decrease in gene expression level was observed even when excluding 300 genes that were expressed at a very low level (TPM <1). Decreased levels of expression in genes with somatic L1 insertions may be due in part to the NMD process triggered by PTC-containing transcripts with abnormal splicing, as suggested above regarding *MOV10*.

## DISCUSSION

It remains an open question whether transposable elements, particularly L1s, play a role in tumorigenesis and what factors determine variable L1 retrotransposition rates in tumors. Here, we analyzed ~200 cancer genomes from three types of gastrointestinal cancer samples and identified a large set of somatic L1 insertions. We found the insertions in some known cancer genes, including *LRP1B*, *PTPRT, ROBO1*, and *PARK2,* in multiple cancer samples. We performed an integrative analysis using RNA-seq profiles of these samples, and found that ~35% of somatic L1 insertions in genes were expressed at a moderate to high level and that they generally disrupted expression of target genes. We also detected somatic retrotransposon insertions causing splicing aberrations.

Although the number of somatic L1 insertions varied greatly in different samples, we detected high average rates of insertions in gastrointestinal cancer, consistent with previous reports. Our analysis identified multiple clinical correlates of somatic L1 retrotransposition in cancer including advanced patient age and tumor stage. *TP53* mutations were frequently observed in cancers with a high rate of retrotransposition, supporting the function of *TP53* as an L1 suppressor. In addition, we discovered a negative correlation between L1 insertion frequency and expression of a subset of known L1 suppressors, including specific *APOBEC3* family proteins and *SMAHD1*, suggesting that they are dominant L1 suppressors in primary human tumors.

Notably, we found that immune activity of tumors is a major factor in explaining the L1 retrotransposition rate. Less L1 retrotransposition activity was found in tumors with high immune activity triggered largely by exogenous or endogenous immunogens such as EBV infection or unstable microsatellite repeats. Moreover, many L1 suppressor genes such as AID/APOBEC, cytidine deaminases, are downstream effector genes of INF and other immune pathways. Although strong immune response effectively suppresses L1 retrotransposition, the increased expression of some APOBECs may lead to extensive APOBEC-mediated mutagenesis, increasing the risk of cancer development (Roberts et al. 2013; Rebhandl et al. 2015). Most importantly, when cancers have low immune activity led by such things as a high CNV load, immune gene aberration, *TP53* mutation, or cancer treatment, they are prone to extensive L1 retrotransposition and thus are at increased risk of tumorigenic insertion events. Larger scale cancer genome analyses considering clinical, genetic, and environmental factors will further illuminate the role of transposable elements in cancer.

## Supplementary Figure legend

**Supplemental Figure S1. Various correlates of somatic L1 retrotransposition.** (a) Cytolytic activity of tumor infiltrating lymphocytes recapitulating cancer immune subgroups. The geometric mean of expression levels of *GZMA* and *PRF1* in each cancer sample was shown with a colored dot. (b) Distinctively high SNV/indel count in high immunity colorectal cancers (CRC-High). (c) Negative correlation between L1 expression and cancer immune activity, notably in stomach/esophageal cancers.

**Supplemental Figure S2. Fraction of differentially expressed immune pathways between the groups**. Among 179 Reactome immune pathways, the percentages of pathways showing up-and down-regulation between immune high *vs* low cancers (FDR < 0.05) is shown for each immune pathway category (innate, adaptive, and cytokine signaling). The numbers of pathways showing up-and down-regulation in high immune subgroups are written in parentheses in red and blue, respectively.

**Supplemental Figure S3. Expression levels of genes with somatic L1 insertions.** The number of genes in each category is shown in parentheses.

## METHODS

### Whole genome sequencing data

We downloaded TCGA whole genome sequencing (WGS) dataset from CGHub https://cghub.ucsc.edu/). The dataset was comprised of bam files for 62 colorectal, 40 stomach, and 19 esophageal cancer samples and matched germline (blood) samples. We downloaded a non-TCGA WGS dataset from EGA (accession id: EGAD00001000782) containing bam files for 55 stomach cancer samples and matched germline samples (Wang et al. 2014). We also downloaded a non-TCGA WGS dataset from dbgap (phs000598.v1.p1) containing bam files for 13 esophageal cancer samples and matched germline samples (Dulak et al. 2013). We realigned the 110 bam files (normal and tumor) in the stomach cancer dataset using the hg19 reference genome and BWA (version 0.6.2) (Li and Durbin 2009). We also marked PCR duplicates for those bam files using picard (http://broadinstitute.github.io/picard). These files do not include bam files that failed to download, realign, or be run with the Tea pipeline, which were excluded from our analysis. Currently the TCGA data is hosted at the Genomic Data Commons (https://gdc.nci.nih.gov/).

### RNA sequencing and gene expression data

We obtained RNA sequencing bam files for 112 TCGA cancer samples from CGHub and gene-level expression data for the TCGA samples from the UCSC cancer genomics browser (https://genome-cancer.ucsc.edu/). For non-TCGA stomach cancer samples, we downloaded raw expression array data from the European Genome-phenome Archive (EGA; accession ID: EGAD00010000528) and extracted gene-level expression data using the IlluminaExpressionFileCreator module in GenePattern (Reich et al. 2006). For non-TCGA esophageal cancer samples, we downloaded gene-level expression data from the Gene Expression Omnibus (GEO; accession ID: GSE42363). We used ComBat (Johnson et al. 2007) to combine expression data from different studies for each cancer type.

### Detection of somatic L1 insertions

We implemented a 3’ transduction calling module in the previously developed transposon detection pipeline (Lee et al. 2012), the transposable element analyzer (Tea), and used this improved version to identify somatic L1 insertions. Each insertion was classified as one of the three types of events defined by Tubio et al (Tubio et al. 2014): solo, partnered, or orphan events. In order to be identified as an insertion, the insertion candidate must have had a poly-A tail and a target-site duplication (TSD) ranging from 5 bp to 35 bp long.

#### Partnered transduction

To detect an L1 insertion with 3’ transduction, repeat-anchored mate (RAM) clusters obtained from *Tea* were paired with discordant read pair clusters generated by *Meerkat* (Yang et al. 2013). For each discordant read pair cluster, we required each to be supported by at least three discordant read pairs. If a RAM cluster was paired with one of the ends from a discordant read pair cluster in positive and negative orientations and the distance between the ends of the clusters was within 1kb, then the two different clusters were merged to define an initial candidate. The remaining ends from the discordant read pair were used to infer the source of an L1 element. To remove false positives, RAM clusters paired with multiple discordant read pair clusters were filtered out. Contigs were assembled using clipped and discordant reads supporting an insertion event, and the assembled contigs were examined for the presence of poly-A tails and TSD.

#### Orphan transduction

For some L1 retrotransposition events with 3’ transductions, 5’ truncation can occur before the retrotransposition of the L1 sequence itself starts, leaving the insertion site with only the 3’ transduction sequence but not the L1 sequence. This type of L1 insertion is defined as an orphan transduction (Tubio et al. 2014). To detect such an event, we used discordant read pair clusters generated by *Meerkat* (Yang et al. 2013). Discordant read pair clusters supported by at least three reads were considered. Two different discordant clusters mapped within 1kb that support both sides of an insertion in the expected orientation were merged to define initial candidates. Clusters mapped with multiple clusters were filtered out. Contigs were assembled using clipped and discordant reads supporting an insertion candidate. The assembled contigs were examined for the presence of poly-A tails and TSD. We called an insertion candidate as an orphan transduction event when its source L1 element was identified as a reference L1HS-Ta, a non-reference germline L1, or a somatic L1 insertion.

### Somatic SNV/indel and copy number aberration call sets

We generated somatic SNV and indel call sets for TCGA colorectal cancer samples using Mutect (Cibulskis et al. 2013) and Varscan2 (Koboldt et al. 2012) with default options, respectively. We annotated the mutations using Oncotator (Ramos et al. 2015). For TCGA stomach and esophageal cancer samples, we obtained somatic SNV/indel call sets from the UCSC cancer genomics browser. For non-TCGA datasets, we obtained somatic SNV/indel call sets from supplementary data in original publications. Only non-synonymous mutations were used in the analyses. We observed that among the TCGA colorectal cancer samples for which clinical MSI assays were performed, all the cases with the MSI-High phenotype except two cases carried more than 100 non-synonymous indels whereas all the non-MSI-High cases carried less than 100 non-synonymous indels. Thus, we classified cancer samples with more than 100 non-synonynous indels as MSI-High (n=2). Somatic copy number aberration data were downloaded from the UCSC cancer genomics browser. The data provided gene-level copy number changes estimated using the GISTIC method (Mermel et al. 2011).

### L1 expression quantification

Reads from RNA-sequencing data were aligned to an L1 sequence library using BWA (Li and Durbin 2009). The L1 sequence library included the L1Hs consensus sequence in Repbase (http://www.girinst.org/repbase/) and its variants created by diagnostic nucleotide substitutions for Ta-1d, Ta-1nd_G1, Ta-1n_C, Ta-0, and Pre-Ta_ACG_G subfamilies. It also included L1Hs sequences that were > 6 kb in size and with a divergence score (relative to the consensus) < 5% in the human reference genome (hg19) annotated by RepeatMasker (http://www.repeatmasker.org/). Reads mapping to Alu and SVA sequence library were excluded. The number of reads mapping to the L1 sequence was normalized by the total number of RNA-seq reads.

### Gene expression analysis

For each gene with a somatic L1 insertion, we calculated the difference between the expression level of the gene from the cancer sample with the insertion and the average expression level from cancer samples without any mutation in the gene. We then calculated the average of the expression differences for each cancer type. To calculate the P value of the observed average expression difference, we estimated a background distribution by using 10,000 randomly selected gene sets with the same number of genes as in our gene set having somatic L1 insertions for each cancer type. The empirical P value was calculated as the proportion of the random gene sets that produced an average expression difference that was less than the observed value.

### Pathway activity analysis

We obtained a set of pathways from the Reactome database (Fabregat et al. 2016). Using ssGSEA (Barbie et al. 2009) in GenePattern (Reich et al. 2006), we measured the activity of each pathway in each cancer sample based on the expression level of its member genes. We obtained additional gene sets reflecting diverse aspects of the immune system (Rooney et al. 2015) and known L1 inhibitors from previous publications (Goodier 2016). Cancer samples were clustered based on the activities of immune pathways using non-negative matrix factorization (NMF) (Brunet et al. 2004) in GenePattern with default parameters (Reich et al. 2006). We separately performed clustering for STAD/ESO and CRC with k=2.

### Gene set enrichment analysis

We tested if genes with somatic L1 insertions in more than one cancer sample were enriched in certain Gene Ontology (GO) terms in the biological process category. We used Goseq (Young et al. 2010) that allowed for adjustment of gene length bias and took as inputs the longest isoform length (the sum of the lengths of all unique exons and introns) for each gene. To identify DNA repair pathways associated with somatic L1 insertion frequency, we obtained 15 DNA repair pathways from a previous publication (Jeggo et al. 2016) and classified cancer samples into the mutant and wild-type groups depending on the presence of a non-synonymous mutation in any member gene for each pathway. We then compared L1 insertion frequencies between the two groups using the Mann-Whitney test.

### Splicing analysis

To identify somatic L1 insertion-mediated abnormal splicing, we employed our previously established ratio-based approach to detect altered splicing caused by somatic mutations (Jung et al. 2015). Specifically, we first extracted abnormally spliced reads near retrotransposon insertion loci and then calculated the ratio of abnormally spliced reads to total reads (the sum of normally and abnormally spliced reads). Uniquely aligned reads excluding PCR duplicates were subjected to this analysis. Next, we assessed whether the ratio was significantly higher than expected, given the background distribution estimated from normal control samples and tumor samples (up to 111 TCGA tumor samples) that lacked non-synonymous mutations for a given gene. We obtained a total of 2,860 control normal RNA-seq data from the Genotype-Tissue Expression (GTEx) project (Consortium 2013). To confirm that the observed splicing change was caused by a somatic retrotransposon insertion, the observed ratio had to be within the top 1% of the background ratio distribution. We screened 69 additional TCGA cancer RNA-seq data to detect splicing aberration caused by somatic retrotransposon insertions reported in a previous study (Helman et al. 2014).

### Analysis of allelic expression of *MOV10*

We called SNVs in *MOV10* from WGS and RNA-seq data from TCGA cancer samples using HyplotypeCaller (DePristo et al. 2011). We excluded cancer samples when they had any non-synonymous mutation, copy number aberration, or DNA methylation (beta value > 0.3) in *MOV10*. For each heterozygous SNV site, both reference and alternative alleles were required to have at least five reads in WGS data to be included in our analysis. In RNA-sequencing data, at least one of the alleles was required to have five reads. A total of 21 cancer samples with at least five heterozygous SNVs in *MOV10* satisfying the minimum read count requirement were used in the analysis as control samples. We used Fisher’s exact test to identify different allelic ratios between DNA-and RNA-sequencing data for each SNV site. We then combined P values from all heterozygous SNV sites in *MOV10* using Fisher’s method. The combined P value was calculated for the TCGA cancer sample (V5-A7RC) with a somatic L1 insertion in *MOV10* as well as for each of the 21 control cancer samples.

## ACKNOWLEDGMENTS

E.L. was supported by K01AG051791, the Harvard Medical School Eleanor and Miles Shore Fellowship, and the Randolph Hearst Fund. H.J. was supported by a grant from NRF (National Research Foundation of Korea) funded by the Korean Government (NRF-2016H1A2A1907072). We thank Hyojin Kang and Junehawk Lee at Supercomputing Center of the Korea Institute of Science and Technology Information for providing computing resources and technical support. The results published here are in part based upon data generated by The Cancer Genome Atlas managed by the NCI and NHGRI. Information about TCGA can be found at http://cancergenome.nih.gov.

